# An Implementation Intention Combining Reappraisal and Anger Counteraction Reduces Fear Reinstatement

**DOI:** 10.1101/2025.09.26.678788

**Authors:** Hongbo Wang, Yingzhu Zeng, Siwen Zeng

## Abstract

Fear extinction requires exposure to the conditioned stimulus (CS), yet patients with fear- and anxiety-related disorders often avoid the CS, thereby undermining extinction learning. Implementation intention (II) is a self-regulation strategy that automates goal-directed behavior through “if–then” planning. Building on the approach motivation elicited by anger and the principle of emotion–action incompatibility, we designed a novel II: *“If I feel afraid when seeing fish/bird images (the CSs), then I will remind myself that it is foolish to fear a mere picture, and I will clench my fists and feel angry with myself*.*”* This strategy aims to transform fear-driven avoidance into self-directed anger, thereby promoting reappraisal and approach behavior. In a three-day differential fear conditioning paradigm, participants who practiced this II during extinction exhibited shorter avoidance distances to the CS+ than the standard-extinction control group. On Day 3, reinstatement testing revealed significantly lower subjective fear, anger, threat expectancy, and avoidance in the II group. These findings suggest that the combined reappraisal-plus-anger II requires practice during extinction to effectively reduce fear return without inducing overt aggression, offering a promising direction for optimizing exposure-based therapies.

## 1. Introduction

Fear extinction is an adaptive learning process in which a conditioned fear response diminishes following repeated exposure to a conditioned stimulus (CS) in the absence of the unconditioned stimulus (US). Through this process, a new “CS–no US” safety memory is formed, which competes with the original “CS–US” fear memory to regulate fear expression. From a theoretical perspective, extinction provides a fundamental model for understanding how competing associative memories interact to shape emotion regulation. From a clinical perspective, extinction principles underlie exposure therapy, which is a first-line treatment for fear- and anxiety-related disorders. However, despite its central role, exposure therapy is not effective for all patients (Loerinc et al., 2015).

A major clinical challenge is that extinction requires sufficient exposure to the CS, yet patients with fear- and anxiety-related disorders often engage in maladaptive avoidance when fear is overwhelming. Avoidance, a defining feature of these disorders (American Psychiatric Association, 2013), reduces contact with the CS, impairs safety learning, and contributes to extinction resistance and relapse (Lovibond et al., 2009; O’Malley & Waters, 2018; Rattel et al., 2017; Vervliet & Indekeu, 2015). For example, O’Malley and Waters (2018) used eye-tracking to direct participants either toward the CS (monitoring) or away from it (avoidance) during extinction training and found that avoiding the CS impaired extinction outcomes. These findings highlight the theoretical importance of examining mechanisms that can counteract avoidance, and the clinical urgency of developing strategies that directly reduce avoidance while enhancing exposure.

Implementation intention (II) represents one promising tool. Structured as “if… then…” statements (e.g., “If I encounter situation X, then I will do Y”), II establishes an automated link between situational cues and goal-directed responses, reducing reliance on conscious effort (Gollwitzer & Sheeran, 2006, 2025). Previous research shows that II can downregulate negative emotions through avoidance (e.g., “If I see a spider, then I will ignore it”; Gallo et al., 2009), suppression (e.g., “If I see a weapon, then I will stay calm”; Azbel-Jackson et al., 2016), or reappraisal (e.g., “If I see blood, then I will view it as a sign of vitality”; Ma et al., 2019). However, these approaches have not fully exploited the emotion incompatibility/counteraction and have rarely been applied to fear extinction.

According to the incompatible response hypothesis (Baron, 1984) and the mutual promotion and counteraction (MPMC) theory of emotionality (Zhan et al., 2017), eliciting responses that counteract fear-driven avoidance can neutralize fear’s behavioral impact. Fear typically promotes avoidance (flight/freeze), whereas anger promotes approach (resistance/confrontation) (OToole & Mikkelsen, 2021). Despite their shared negative valence, their action tendencies are opposed. This makes anger a potential counterforce to fear-driven avoidance. Physiologically, fear and anger share overlapping threat-related neural substrates (Prather, 2016; Siegel et al., 2018), which makes fear-to-anger transformation feasible (Zhan et al., 2015, 2018). Functionally, anger enhances certainty and perceived control (Lerner & Keltner, 2001; Song et al., 2021; Tiedens & Linton, 2001), fosters a sense of personal power (Sell et al., 2009; Tibubos et al., 2013), and facilitates goal pursuit (Lench et al., 2024). Notably, an individual’s sense of certainty and control serves as a critical factor in mitigating the negative effects of stress (Hartley et al., 2014; Meyer et al., 2021). Historical case reports even document therapeutic effects of anger induction in treating fear symptoms (Butler, 1975; Goldstein et al., 1970). For example, Goldstein et al. (1970) instructed patients to pair fear-evoking scenarios (either imagined or real) with anger imagery, accompanied by vocal and motor expressions, before applying these techniques to real-life fear stimuli—ultimately achieving therapeutic effects. Greenberg & Pascual-Leone (2024) also proposed that the experience of implicit adaptive anger toward the aggressive acts of perpetrators contributes to modifying maladaptive fear in trauma survivors.

Yet, directly inducing anger raises ethical and clinical concerns. On one hand, anger is a symptom in some anxiety and trauma disorders. For instance, the DSM-5 diagnostic criteria for PTSD include anger within Criterion D (“Negative Alterations in Cognitions and Mood”) and Criterion E (“Alterations in Arousal and Reactivity”), specifically “irritable behavior and anger outbursts” (American Psychiatric Association, 2013). On the other hand, anger can contribute to aggression (Chereji et al., 2012). Further analysis suggests that within anxiety disorders, it is difficult to distinguish primary from secondary anger and irritability symptoms. For some patients, anger may be a secondary symptom—reflecting “defensive displacement” in response to an uncontrollable threat. This involves redirecting suppressed frustration from a high-threat object (e.g., a social situation) that cannot be directly confronted to a lower-threat object (e.g., family members or other stimuli) (Marcus-Newhall et al., 2000), analogous to the “kicking the cat” effect. For instance, research has indicated that fear of COVID-19 might manifest as hidden aggression within the virtual social ecosystem (Ye et al., 2021). Neurobiological mechanisms suggest that this emotional transfer in anxious individuals may stem from hypervigilance to threat (e.g., amygdala activation) coupled with insufficient prefrontal regulation (Shackman et al., 2011). More importantly, cognitive behavioral therapy for individuals with generalized anxiety disorder, while not directly targeting anger, has been shown to improve both internally experienced and externally expressed anger (Laposa & Fracalanza, 2019). This improvement might occur because treatment attenuates the emotional response to the original threat, reducing the need to vent unresolved anxiety onto “safe targets”—suggesting that such anger symptoms may be a byproduct of unresolved fear/avoidance rather than primary emotional dysregulation. Furthermore, anger does not inevitably lead to aggression; it exists on a continuum (from mild annoyance to intense anger to rage) (Deffenbacher et al., 1996). Forbes et al. (2008) demonstrated that it is specifically “fear of anger” (e.g., “worry about entering an endless state of anger and its harmful consequences”) that predicts poor treatment outcomes.

Nonetheless, to minimize the risks associated with anger induction, we developed a novel II strategy integrating reappraisal and self-directed anger: *“If I feel afraid when seeing fish/bird images (the CSs), then I will remind myself that it is foolish to fear a mere animal picture, and I will clench my fists and feel angry with myself*.*”* This design incorporates three safeguards: (1) pre-emptive reappraisal (establishing “CS = safe” before anger is triggered), (2) conditional activation (anger only arises if fear is detected), and (3) self-focused anger (directed at one’s own irrational fear, not external targets). These features minimize risks while leveraging anger’s approach motivation to counter fear-driven avoidance. Through this integrated “reappraisal + anger counteraction” II, we aim to achieve two primary objectives (see Fig. **1**): first, to prompt a reappraisal of the CS’s threat value, facilitating an automatic approach response; and second, to harness the proactive energy of anger to enhance the willingness to approach the CS, thereby improving extinction efficacy. This design not only addresses ethical concerns but also aligns with the safety requirements for clinical translation.

**Fig. 1.**
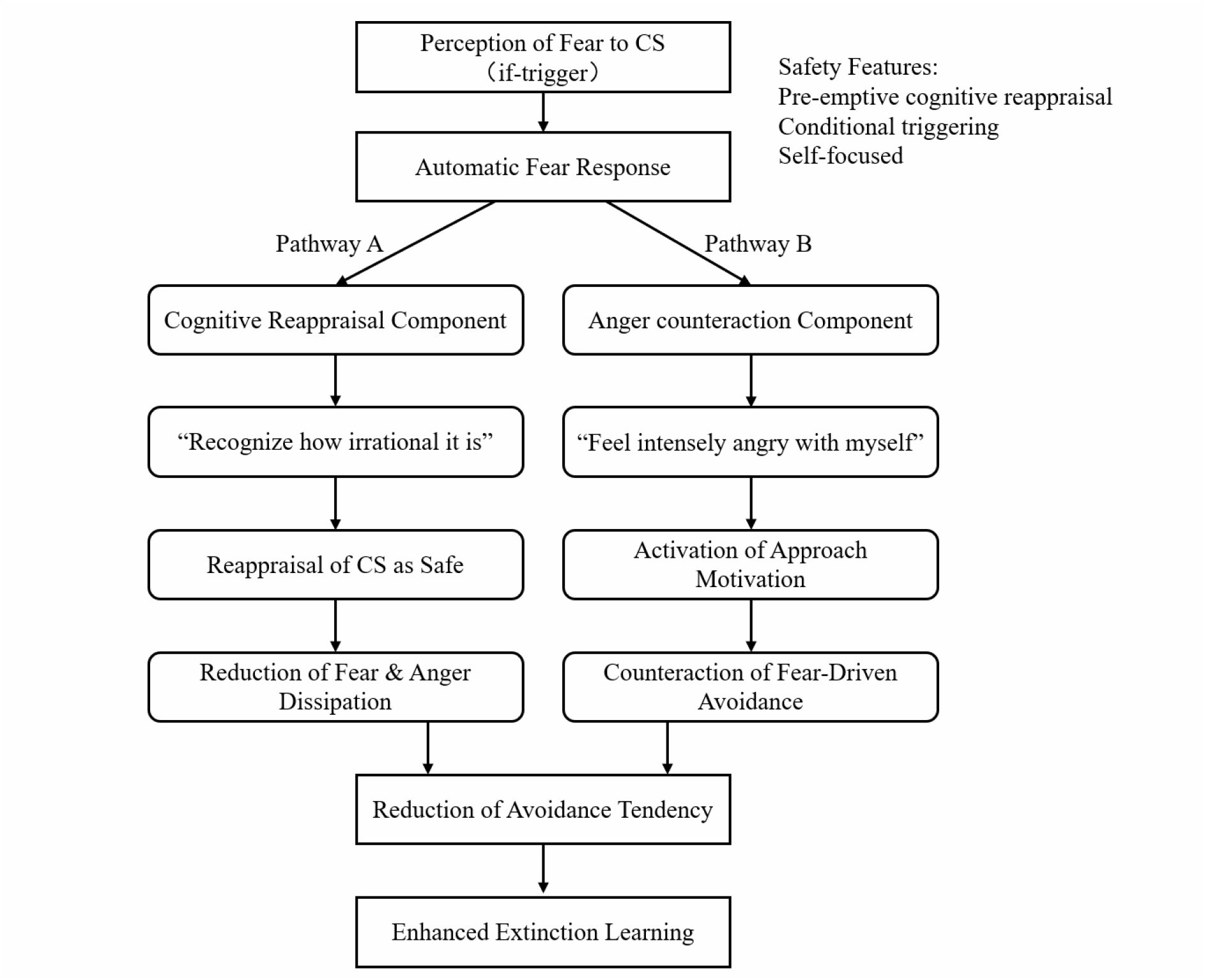
Proposed mechanisms of anger-countering implementation intentions (IIs). Safety features mitigate potential risks associated with anger induction. Pathway A depicts cognitive reappraisal as the dominant mechanism. Pathway B illustrates that emotion counteraction via anger serves to engage the approach motivation.

Based on the incompatible response hypothesis, the automated regulatory function of II, and the approach-motivational characteristics of anger, we hypothesized that, compared with standard extinction, participants who practiced this II would show stronger approach motivation (less avoidance), more consolidated safety learning, and reduced return of fear.

## 2. Methods

### 2.1 Participants

The study compared an implementation intention (II) group with a control (Ctrl) group, using repeated measurements across at least two time points. Sample size was estimated with G*Power 3.1 (Faul et al., 2007), assuming a medium effect size (f = 0.25), α = 0.05, and power (1 – β) = 0.80. With an assumed within-subject correlation of 0.50, the required sample was 34 participants. Considering the high attrition rates typical of fear-conditioning studies, 49 undergraduates (aged 18–25 years) were recruited via advertisements and voluntary sign-ups.

Inclusion criteria were: right-handedness, normal or corrected-to-normal vision, no history of physical or psychiatric disorders, and no prior participation in similar experiments. Participants attended the study across three consecutive days at approximately the same time of day. The study was approved by the Ethics Committee of the School of Psychology, Henan University (approval no. 20210923002). Written informed consent was obtained. Participants were informed that: (1) shock intensity would be individually calibrated within a safe range; (2) they could withdraw at any time; (3) personal data would remain confidential; and (4) they were required to maintain confidentiality about the study procedures. All participants received monetary compensation.

Participants were randomly assigned to the Ctrl group (n = 25) or the II group (n = 24). Data from 12 participants were excluded: three due to failure in category learning on Day 1 (Ctrl: 11, 14; II: 1), eight due to withdrawal because of shock intolerance or discomfort (Ctrl: 15, 18, 19, 22; II: 18–20, 24), and one due to equipment malfunction during Day 3 reinstatement (Ctrl: 23). The final sample included 37 participants (Ctrl: n = 18; II: n = 19). Groups did not differ significantly in age, gender, trait anxiety, depression, or shock intensity (Table **1**).

**Table 1.** Descriptive Statistics for Group Demographics and Questionnaire Scores (M ± SD).

| | II ($n = 19$) | Ctrl ($n = 18$) | $\chi^2 / t$ | $p$ | $d$ |
| --- | --- | --- | --- | --- | --- |
| Male: female | 8:11 | 7:11 | 0.04 | 0.842 |  |
| Age (years old) | 20.74 ± 1.91 | 20.78 ± 2.02 | -0.063 | 0.950 | 0.02 |
| STAI-S | 42.95 ± 4.64 | 44.56 ± 4.22 | -1.102 | 0.278 | 0.363 |
| STAI-T | 45.74 ± 4.70 | 44.00 ± 3.63 | 1.253 | 0.219 | 0.414 |
| STAEI | 37.21 ± 7.08 | 37.33 ± 8.16 | -0.049 | 0.961 | 0.016 |
| IUS | 19.84 ± 4.71 | 20.94 ± 4.61 | -0.719 | 0.477 | 0.236 |
| Shock intensity | 52.40 ± 4.75 | 54.71 ± 10.62 | -0.18 | 0.859 | 0.281 |
| Shock discomfort | 7.53 ± 0.77 | 7.50 ± 0.79 | 0.103 | 0.919 | 0.038 |

Note: Data are presented as mean (standard deviation). II = the implementation intention (II) group; Ctrl = the control (Ctrl) group; STAI-S = State Anxiety Inventory; STAI-T = Trait Anxiety Inventory; STAEI = State Trait Anger Expression Inventory; BDI = Beck Depression Inventory; IUS = Intolerance of Uncertainty Scale.

### 2.2 Stimuli

Following Kroes et al. (2017), conditioned stimuli (CSs) were pictures of fish and birds (148 total) obtained from Pixabay and Baidu. One exemplar from each category was used for explicit CS ratings; the remaining 73 fish and 73 bird images were used for training and testing. All pictures (1280 × 720 px, 96 dpi, 24-bit) were presented for 5 s on a 21-inch LCD monitor.

For one category, 67% of images were followed by an electric shock (CS+), while the other category was never reinforced (CS–). Category assignment (fish vs. bird) was counterbalanced across participants. The unconditioned stimulus (US) was a 200-ms shock delivered to the left wrist via a constant-voltage stimulator (STM200-1, BIOPAC Systems, Inc.). Shock intensity was individually calibrated: participants rated sample shocks on a 1–9 scale (1 = no discomfort, 8 = slight but acceptable pain, 9 = unbearable), and the intensity corresponding to a rating of 8 was used.

### 2.3 Subjective assessments

Subjective ratings for CS+ and CS− were collected (using bird pictures as an example, see Fig. **1D**) as follows: (1) US Expectancy (“How likely is this image to be followed by a shock?” 1 = certainly not, 9 = certainly will) served as a prospective measure of associative memory. (2) CS–US Contingency (“Based on previous learning, how often was a bird picture followed by shock?” 1 = never, 9 = always) served as a retrospective measure of associative memory. (3) Fear level (“How fearful do you feel when seeing the bird picture?” 1 = not at all, 9 = extremely). (4) Anger level (“How angry do you feel when seeing the bird picture?” 1 = not at all, 9 = extremely). (5) Avoidance distance (“How far would you like to be from this picture?” 1 = closest, 9 = farthest).

US expectancy ratings were collected immediately before CS presentation. In contrast, other ratings were obtained pre- and post-block. For avoidance distance, participants moved a blue avatar along a 1– 9 scale to indicate their desired distance from the CS.

Shock discomfort was assessed twice (end of Day 1 acquisition and after Day 3 reinstatement). After the reinstatement test, participants also recalled the number of shocks associated with each CS type.

### 2.4 Physiological Recording

Skin conductance response (SCR) was recorded using two standard Ag/AgCl electrodes (8 mm diameter) filled with 0.05 M sodium chloride electrolyte paste. The electrodes were attached to the palmar surfaces of the index and middle fingers of the participant’s non-dominant hand and connected to an EDA100C amplifier. SCRs were acquired using a Biopac data acquisition system (Model MP150) with a sampling frequency of 2000 Hz running AcqKnowledge software Version 4.4 (Biopac Systems, Inc., Goleta, California, USA). SCR was collected during acquisition (Day 1) and reinstatement (Day 3), but not extinction (Day 2), as fear and anger both activate sympathetic arousal, making differentiation difficult (Damasio & Carvalho, 2013; Stemmler et al., 2001).

SCR was low-pass filtered at 1 Hz (Stussi et al., 2019). SCR for each CS was calculated as the peak-to-peak skin conductance response within a 0.5-5.5 s window post-stimulus onset, measured in microsiemens (μS). To normalize the data distribution, raw SCR values were square-root transformed and standardized by dividing by the participant’s maximum SCR value on the same day (Scheuermann et al., 2025).

### 2.5 Procedure

The experiment was programmed and run using E-Prime 2.0 software (Psychology Software Tools Ltd., Pittsburgh, PA). The procedure spanned three consecutive days: Fear Acquisition (Day 1), Fear Extinction (Day 2), and Spontaneous Recovery/Reinstatement Test plus a Surprise Recognition Memory Test (Day 3). The types of stimuli, stimulus presentation times, and inter-stimulus intervals were consistent across days. The experimental design and timeline are illustrated in Fig. **2**.

**Fig. 2.**
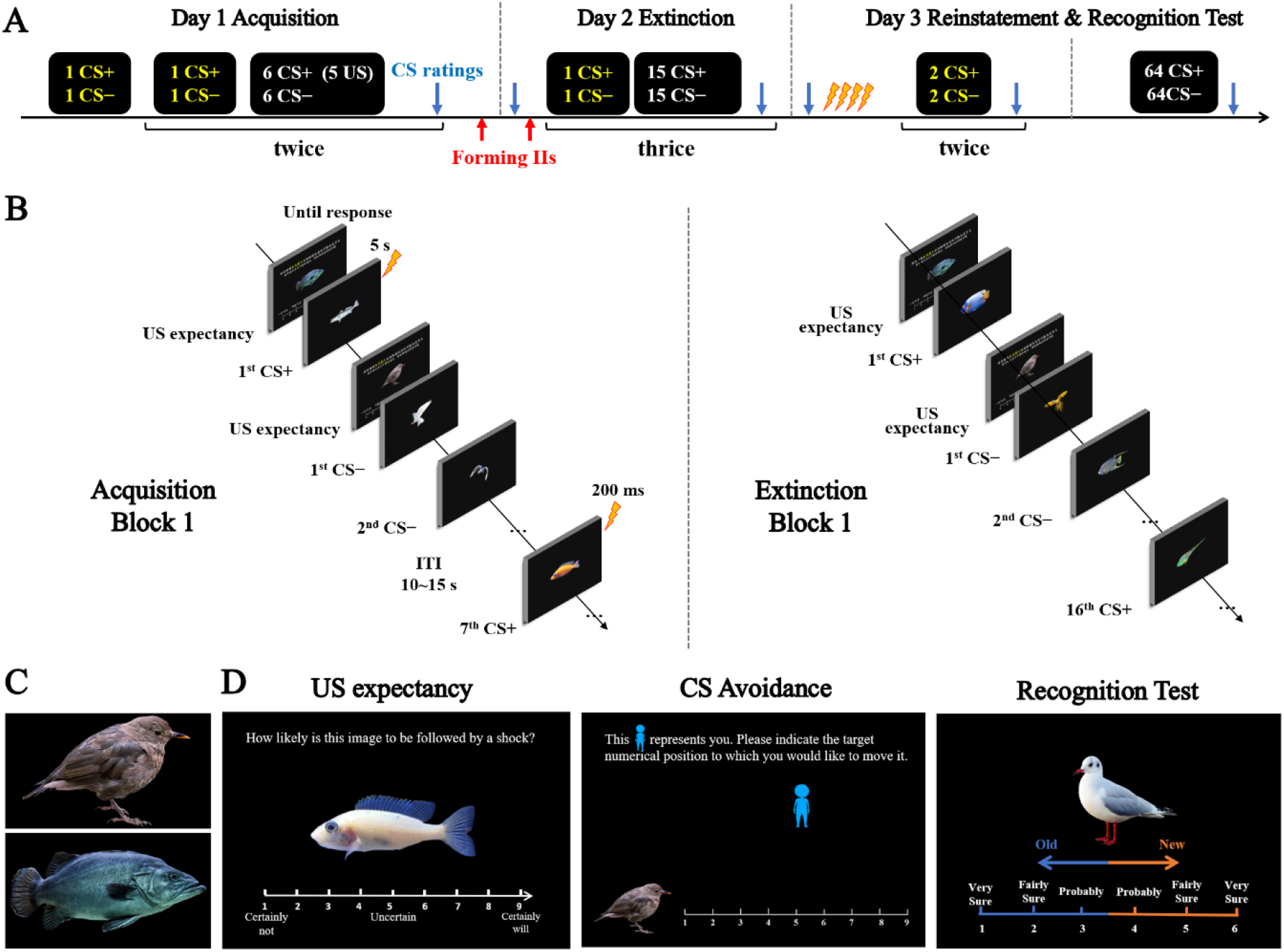
Schematic of Experimental Design and Procedure. (**A**) Schematic of experimental flow and trial sequence. CS markers in yellow indicate trials preceded by a US expectancy rating; white CS markers indicate trials without a preceding US expectancy rating. Blue arrows denote time points at which the four CS-related subjective ratings were collected, including CS–shock contingency, fear level to CS, anger level to CS, and avoidance distance to CS. The red arrow indicates the implementation intention (II) training session conducted for the II group. (**B**) Example trial sequence for Block 1 during fear acquisition on Day 1 (left) and fear extinction on Day 2 (right), using fish pictures as the CS+. “…” indicates omitted trials in the sequence. (**C**) Representative CS images used in the subjective rating interface. (**D**) Illustration of the rating interfaces: US expectancy rating (left), CS avoidance distance rating (middle), and image recognition test (right). Participants responded using the numeric keys according to on-screen instructions.

#### Fear Acquisition (Day 1)

The experimenter explained the procedures, after which participants provided written informed consent. Participants then completed the State-Trait Anxiety Inventory (STAI; Spielberger et al., 1983), Beck Depression Inventory (BDI; A. T. Beck et al., 1996), Trait Anger Scale (Tibubos et al., 2020), and Intolerance of Uncertainty Scale (IUS; Buhr & Dugas, 2002). Subsequently, electrodes for SCR and the shock stimulator were attached. Before the formal experiment, individual shock intensity was calibrated (range: 30-70 V) as described in Section 2.2. This calibrated intensity (corresponding to a rating of 8) was used for all subsequent sessions. After a 4-5 minute rest, instructions were presented. Participants were informed that pictures of fish and birds would be shown and that they should learn and predict the likelihood of a shock following each picture type. The acquisition phase used 15 unique pictures from each category (fish, bird), with a reinforcement rate of 2/3 for the CS+. Specifically, the 1st, 3rd, 5th, 10th, and 12th presentations of the CS+ were not reinforced (pseudorandom sequence). The first two trials consisted of one CS− and one non-reinforced CS+ (order balanced), each preceded by a US expectancy rating. The remaining trials were divided into two blocks of 14 trials each (5 CS+ paired with shock, 2 CS+ not paired with shock, 7 CS−). The first two trials of each block were a CS− and a CS+-noUS trial, both with US expectancy ratings; the remaining 12 trials within the block did not include US expectancy ratings (Fig. **2A, B**). CS presentation order was pseudorandomized within blocks. After each block, participants were prompted to recall which picture category was followed by the shock, followed by the set of four CS-related subjective assessments (CS-US contingency, fear, anger, avoidance distance). A 1-2 minute rest period followed each block. Subjective shock discomfort was rated again at the end of acquisition. Participants in the control group completed the experimental session for Day 1, while those in Group II were required to stay for an additional 3-minute II training. They were instructed to silently recite the following II phrase: *“If I feel afraid when seeing fish/bird images, then I will remind myself that it is foolish to fear a mere picture, and I will clench my fists and feel angry with myself.”* They spent two minutes with their eyes closed, vividly imagining themselves executing this plan. Participants were provided with standardized guiding scripts (e.g., ‘Imagine seeing a fish picture, feeling scared, then thinking “I’m foolish to fear this harmless image” and clenching your fists to feel angry at yourself’) to ensure consistent imagination content across individuals. They were also told that this practice was crucial for the next day’s experiment and that they should rehearse it mentally. This preliminary II training was implemented due to concerns that the training immediately preceding extinction on Day 2 might be insufficient to automatize the II response.

#### Fear Extinction (Day 2)

After sitting quietly for 4-5 minutes, all participants completed the set of four CS-related subjective assessments to check for any baseline differences between groups following the previous day’s II training. Participants in the Control Group proceeded directly to the extinction phase. Participants in the II Group were first asked to recall and type the II phrase from memory. The correct II phrase was then displayed on the screen, and participants rated the similarity between their recollection and the displayed phrase on a scale. If the self-rated similarity was ≥5, they performed 3 minutes of II training; if it was <5, they performed 6 minutes. Only then did the II Group proceed to extinction. The extinction phase used 24 novel pictures from each category (different from Day 1), each presented twice, resulting in 96 trials divided into 3 blocks. Each block contained 16 CS+ and 16 CS− trials presented in a pseudorandom order, none of which were reinforced with shock (Figure 1A, B). As in acquisition, the first two trials of each block consisted of one CS− and one CS+ (order balanced), each preceded by a US expectancy rating; the remaining 30 trials in the block did not include US expectancy ratings. At the end of each block, the four CS-related subjective assessments were administered, followed by a 1-2 minute rest before the next block.

#### Spontaneous Recovery and Reinstatement Test (Day 3)

After sitting quietly for 4-5 minutes, all participants first completed the set of four CS-related subjective assessments to measure spontaneous recovery (SR). Subsequently, participants received four unsignaled US presentations (shocks, 200 ms duration) with inter-stimulus intervals of 20, 30, 25, and 15 seconds. After a 10-minute rest, the reinstatement test began. The reinstatement test consisted of 9 trials. Each trial began with a US expectancy rating, followed by the presentation of a CS picture. The pictures used were 9 “old” pictures from Day 1 (5 CS−, 4 CS+ that had been paired with shock on Day 1). The first trial was always a CS−, followed by two mini-blocks. Each mini-block contained 2 CS+ and 2 CS− trials (order randomized within the mini-block), none reinforced. The set of four CS-related subjective assessments was administered after each mini-block; there was no rest between mini-blocks. Subjective shock discomfort was rated again after the reinstatement test. Participants were also asked to recall and input the number of shocks that had followed each CS type (fish/bird) on Day 1. The limited number of trials in the reinstatement test was based on the assumption that group differences would be most pronounced in the initial trials, after which extinction would quickly occur in both groups due to the extensive extinction training (96 trials) on Day 2. Furthermore, potential fear renewal after reinstatement was not a major concern, as the subsequent surprise recognition memory test involving 136 CS presentations was expected to serve as additional extinction.

#### Surprise Recognition Memory Test (Day 3)

Following the reinstatement test, participants performed an unexpected recognition memory test. They were asked to judge whether each presented picture was “old” (seen during Days 1 or 2) or “new” (not seen before). This test aimed to investigate whether the anger-based II strategy operated automatically without increasing cognitive effort. For “old” responses, participants pressed keys 1, 2, or 3, with lower numbers indicating higher confidence in having seen the picture. For “new” responses, participants pressed keys 4, 5, or 6, with higher numbers indicating higher confidence in not having seen the picture. The test comprised 136 pictures (68 fish, 68 birds), including 10 old pictures from Day 1 (excluding those used in the reinstatement test), 24 old pictures from Day 2, and 34 new pictures. Pictures were divided into 6 groups based on their novelty (Day 1 old, Day 2 old, new) and CS type (CS+, CS−). The order of these groups was pseudorandomized to prevent more than two consecutive pictures from the same group, although the order of pictures within each group was random. During each trial, a picture was presented with a 1-6 confidence scale displayed below it. The picture disappeared upon key press. The inter-trial interval (ITI) was a red fixation cross presented for 1 second. After the memory test, the set of four CS-related subjective assessments was administered once more. The experiment concluded, and participants were debriefed and compensated. The post-reinstatement recognition test procedure was adapted from Kroes et al. (2017).

### 2.6 Data analyses

All data analyses were conducted using SPSS 26.0. Continuous variables are presented as means ± standard deviations (M ± SD). Between-group baseline differences (e.g., in age and trait anxiety) were assessed via chi-square (χ^2^) tests or Welch-corrected independent t-tests. Subjective assessment data (CS-US contingency, fear, anger, avoidance distance, US expectancy) from the fear acquisition, extinction, spontaneous recovery, and reinstatement phases were analyzed using repeated-measures ANOVAs with the dependent variable being the CS difference (CS+ minus CS−), to evaluate the interaction and main effects of Group (II, Ctrl) × Test Time (prior to or after blocks, or test time points). SCR and recognition test data were analyzed via two-way repeated-measures ANOVAs with Group (II, Ctrl) and CS type (CS+, CS−) as factors. Significant interactions were followed by simple effects analyses with Bonferroni corrections for multiple comparisons. Effect sizes are reported as partial η^2^. The significance level for all statistical tests was set at α = 0.05 (two-tailed).

## 3. Results

### 3.1 Preliminary Analyses

No significant group differences were observed in sex ratio, age, state or trait anxiety, trait anger, depression, intolerance of uncertainty, shock intensity, or shock discomfort (see Table **1**).

### 3.2 Subjective Assessments

CS rating differences (CS+ minus CS−) for four CS-related subjective assessments (CS–US contingency, fear, anger and avoidance distance) and US expectancy across the fear acquisition, extinction, spontaneous recovery, and reinstatement phases are shown in Fig. **3**.

**Fig. 3.**
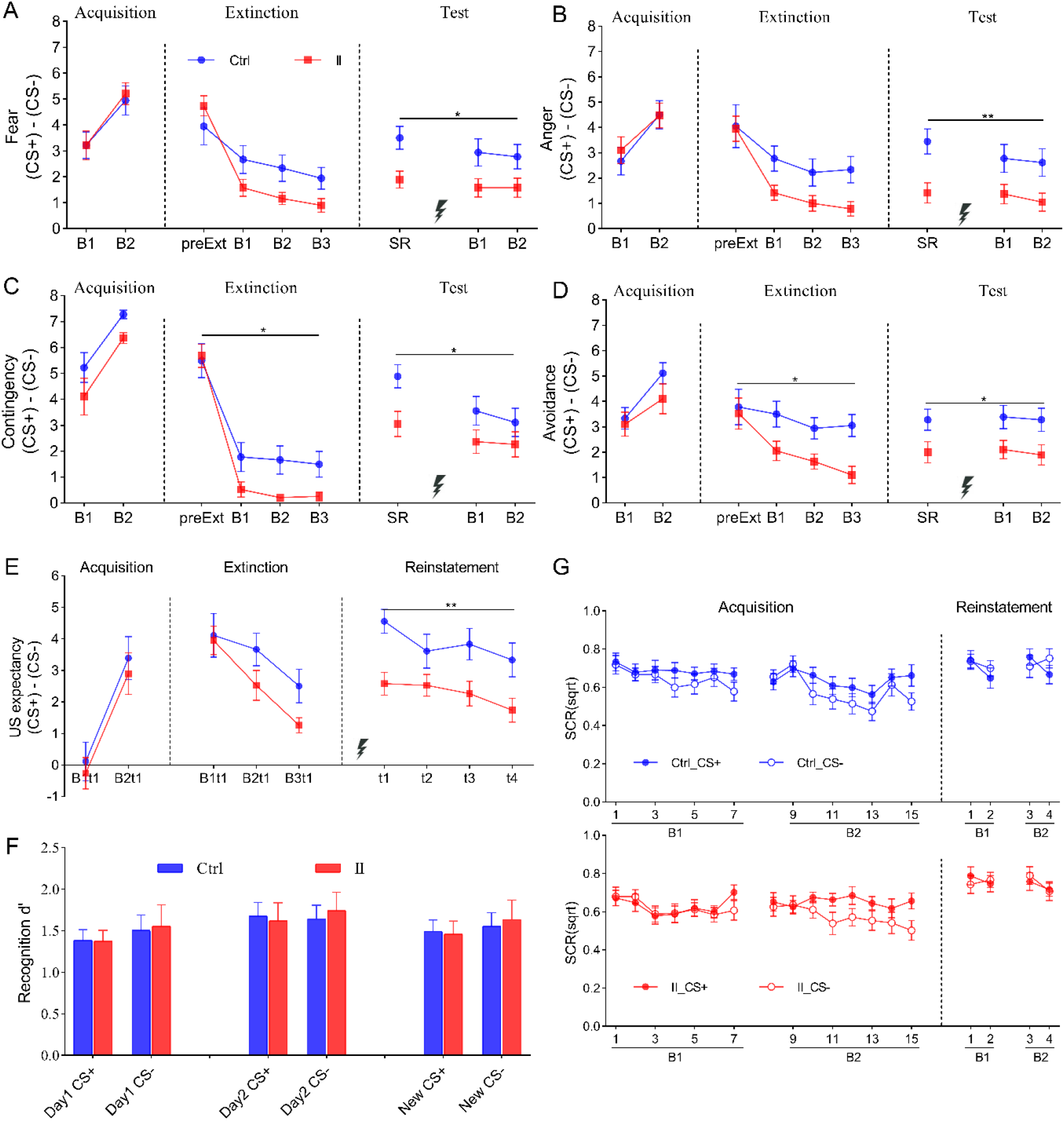
Subjective ratings, SCRs, and recognition memory performance. Data shown are difference scores (CS+ minus CS−) for subjective measures and SCRs across phases, and recognition test results on Day 3 for the implementation intention (II) and control (Ctrl) groups. (**A**) Fear ratings. (**B**) Anger ratings. (**C**) CS–US contingency ratings. (**D**) Avoidance distance. (**E**) US expectancy. (**F**) Recognition memory performance. (**G**) SCR data during fear acquisition and reinstatement. Abbreviations: B1/B2/B3, post-Block 1/2/3; preExt, pre-extinction on Day 2; SR, spontaneous recovery test; B1t1/B2t1, the first trial of Block 1/2; t1–t4, Trials 1–4 of reinstatement.

#### Fear acquisition (Day 1)

Two-way repeated-measures ANOVAs with Group (II, Ctrl) and Test Time (B1, B2) as factors was conducted on the CS differences (CS+ minus CS−) for four CS-related subjective assessments assessed post Block 1 (B1) and Block 2 (B2). For US expectancy assessed during the first trial of each block, a two-way repeated-measures ANOVA with Group (II, Ctrl) and Test Time (B1t1, B2t1) as factors was conducted on the CS differences (CS+ minus CS−). Results indicated no significant main effects of Group (fear: *p* = 0.841; anger: *p* = 0.769; contingency: *p* = 0.06; avoidance: *p* = 0.336; US expectancy: *p* = 0.458). The main effects of Test Time were significant, with all assessments showing larger CS differences at B2t1 than B1t1 (fear: *F*(1, 35) = 25.524, *p* < 0.001,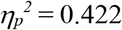 anger: *F*(1, 35) = 30.448, *p* < 0.001, 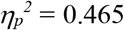 contingency: *F*(1, 35) = 24.466, *p* < 0.001, 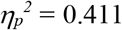 avoidance: *F*(1, 35) = 28.013, *p* < 0.001, 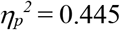 US expectancy: *F*(1, 35) = 24.356, *p* < 0.001, 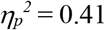). No significant Group × Test Time interactions were found (fear: *F*(1, 35) = 0.142, *p* = 0.708; anger: *F*(1, 35) = 0.642, *p* = 0.428; contingency: *F*(1, 35) = 0.057, *p* = 0.813; avoidance: *F*(1, 35) = 2.196, *p* = 0.147; US expectancy: *F*(1, 35) = 0.008, *p* = 0.927). These results indicate significant changes in participant responses across acquisition phases, with all assessments showing increased CS differences and successful conditioning of differential fear responses in both groups.

### Fear extinction (Day 2)

Two-way repeated-measures ANOVAs with Group (II, Ctrl) and Test Time (preExt, B1, B2, B3) as factors were performed on the CS differences (CS+ minus CS−) for four CS-related subjective assessments assessed prior to the first extinction block (preExt) and post each extinction block (B1/B2/B3). Results showed no significant main effects of Group for fear (*p* = 0.218) or anger (*p* = 0.075). However, the Ctrl group exhibited significantly higher contingency (*F*(1, 35) = 5.218, *p* = 0.029, 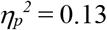) and avoidance (*F*(1, 35) = 5.133, *p* = 0.03, 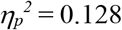) than the II group, indicating that the IIs facilitated fear extinction and reduced avoidance of CS+. The main effect of Test Time was significant (fear: *F*(3, 105) = 38.151, *p* < 0.001,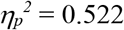 anger: *F*(3, 105) = 24.055, *p* < 0.001, 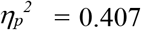 contingency: *F*(3, 105) = 69.916, *p* < .001,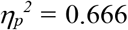 avoidance: *F*(3, 105) = 9.415, *p* < 0.001, 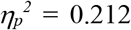). Bonferroni post-hoc tests revealed that at the end of extinction (B3), the CS differences for all four assessments were significantly lower than preExt (fear: *p* < 0.001; anger: *p* < 0.001; contingency: *p* < 0.001; avoidance: *p* = 0.007). No significant Group × Test Time interactions were found for anger, contingency, or avoidance (anger: *p* = 0.164; contingency: *p* = 0.165; avoidance: *p* = 0.100). However, a significant Group × Test Time interaction was observed for fear (*F*(3, 105) = 4.933, *p* = 0.012, 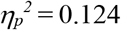). Simple effects analysis indicated no inter-group differences in fear scores at preExt and B1 (preExt: *p* = 0.328; B1: *p* = 0.088), but significant differences at B2 and B3 (B2: *p* = 0.038; B3: *p* = 0.038). In the II group, fear scores at preExt were significantly higher than B1, B2, and B3 (all *p’s* < 0.001), with no differences among the latter three time points (B1 vs. B2: *p* = 0.861; B1 vs. B3: *p* = 0.218; B2 vs. B3: *p* = 1.000). In the Ctrl group, fear scores at preExt were significantly higher than B2 and B3 (B2: *p* = 0.032; B3: *p* = 0.007), with no differences among B1, B2, and B3 (B1 vs. B2: *p* = 1.000; B1 vs. B3: *p* = 0.191; B2 vs. B3: *p* = 0.899). This suggests that both groups showed fear expression at the start of Day 2, but after a phase of extinction training, fear expression toward CS+ decreased significantly, with the II group exhibiting superior extinction outcomes compared to the Ctrl group.

A two-way repeated-measures ANOVA was conducted on CS differences (CS+ minus CS−) for US expectancy during extinction, with Group (II, Ctrl) and Test Time (B1t1, B2t2, B3t3) as factors. Results indicated no significant main effect of Group (*p* = 0.149) or Group × Test Time interaction (*p* = 0.251). However, the main effect of Test Time was significant (*F*(2, 70) = 18.542, *p* < 0.001, 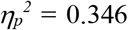). Bonferroni post-hoc tests revealed that US expectancy differences in the first trial of Block 3 (B3t1) were significantly lower than B1t1 and B2t1 (B1t1 vs. B3t1: *p* < 0.001, B2t1 vs. B3t1: *p* = 0.001, B1t1 vs. B2t1: *p* = 0.035). This indicates that by the end of extinction, both groups showed a marked decrease in US expectancy for CS+, consistent with successful extinction and aligning with other assessment metrics.

#### Fear recovery and reinstatement (Day 3)

Two-way repeated-measures ANOVAs with Group (II, Ctrl) and Test Time (SR, B1, B2) as factors were conducted on the CS differences (CS+ minus CS−) for four CS-related subjective assessments assessed during the spontaneous recovery (SR) phase before reinstatement and after each reinstatement test block (B1/B2). The II group showed significantly lower scores across all four assessments than the Ctrl group (fear: *F*(1, 35) = 6.54, *p* = 0.015,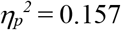 anger: *F*(1, 35) = 7.787, *p* = 0.008, 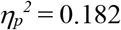 contingency: *F*(1, 35) = 4.59, *p* = 0.039, 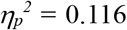 avoidance: *F*(1, 35) = 6.000, *p* = 0.019, 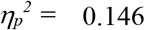). Significant main effects of Test Time were found for fear (*F*(2, 70) = 3.661, *p* = 0.039, 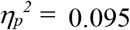) and contingency (*F*(2, 70) = 7.918, *p* = 0.004, 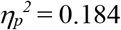), with a marginal effect for anger (*F*(2, 70) = 3.65, *p* = 0.052, 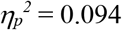). No significant main effect of Test Time was found for avoidance (*p* = 0.747). Bonferroni post-hoc tests revealed that during the spontaneous recovery (SR) phase before reinstatement, CS differences for fear and contingency were significantly higher than post-reinstatement (B2) (both *p’s* < 0.05), with no differences between other time points. No significant Group × Test Time interactions were observed (fear: *p* = 0.58; anger: *p* = 0.338; contingency: *p* = 0.282; avoidance: *p* = 0.924).

Following US reinstatement, a two-way repeated-measures ANOVA was performed on the CS differences (CS+ minus CS−) for US expectancy across four trials, with Group (II, Ctrl) and Trial (t1, t2, t3, t4) as factors. The analysis revealed a significant main effect of Group (*F*(1, 35) = 7.588, *p* = 0.009, 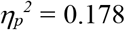), indicating that the Ctrl group had significantly higher US expectancy differences than the II group. A significant main effect of Trial was also observed (*F*(3, 105) = 10.066, *p* < 0.001,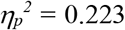). Bonferroni post-hoc tests showed that US expectancy in trial 4 was significantly lower than in trials 1, 2, and 3 (*p* < 0.001, *p* = 0.029, *p* = 0.002, respectively), demonstrating a rapid decline in US expectancy for CS+ across the four trials. The interaction between Group and Trial was not significant (*p* = 0.149). These findings indicate that the IIs strategy effectively reduces conditioned fear return.

### 3.3 Surprise Recognition Test (Day 3)

A two-way ANOVA conducted on the surprise recognition test data with Group (II, Ctrl) and CS type (CS+, CS−) as factors, revealed no significant main effects or interactions (see Fig. **3F**). For old Day 1 images, the main effect of Group was non-significant (*p* = 0.921). Similarly, no significant Group effects were found for old Day 2 images (*p* = 0.935) or new images (*p* = 0.923). No significant main effect of CS type was found (Day 1 old: *p* = 0.25; Day 2 old: *p* = 0.489; new: *p* = 0.148). Additionally, no significant interactions between Group and CS were observed (Day 1 old: *p* = 0.827; Day 2 old: *p* = 0.212; new: *p* = 0.521).

### 3.4 Skin Conductance Response (SCR)

SCR results for the two groups during fear acquisition and reinstatement are shown in Fig. **3G**.

#### Fear acquisition (Day 1)

Data from one participant in the Ctrl group were unavailable. A two-way repeated-measures ANOVA was performed on the SCR data from the final trial of fear acquisition, with Group (II, Ctrl) and CS type (CS+, CS−) as factors. The analysis revealed a significant main effect of CS type (*F*(1, 34) = 15.256, *p* < 0.001,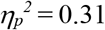), indicating higher SCR values for CS+ compared to CS−. However, no significant main effect of Group (*p* = 0.808) or Group × CS type interaction (*p* = 0.801) was observed.

#### Fear reinstatement (Day 3)

A similar ANOVA on the SCR data from the first trial of reinstatement showed no significant effects. Neither the main effect of CS type (*p* = 0.467), nor the main effect of Group (*p* = 0.647), nor the Group × CS type interaction (*p* = 0.579) reached significance.

## 4. Discussion

This study examined whether forming an implementation intention (II) that combines cognitive reappraisal with self-directed anger could facilitate fear extinction and reduce the return of fear. The results provided converging evidence for this hypothesis. During extinction, participants in the II group reported significantly lower CS–US contingency ratings and showed reduced avoidance distance toward the CS+ compared with the control group. Furthermore, during both the spontaneous recovery and reinstatement tests on Day 3, the II group reported lower levels of fear, anger, contingency, and avoidance distance, while also showing lower US expectancy during reinstatement. Together, these findings suggest that practicing reappraisal-plus-anger IIs during extinction promotes safety learning and provides protection against fear relapse.

However, during the reinstatement test on Day 3, SCR data revealed no significant main effects of Group or CS type, nor a significant Group × CS type interaction. Consistent with prior studies (Haesen & Vervliet, 2015; Scheuermann et al., 2025), subjective ratings and SCRs showed divergent patterns. Several factors may account for the absence of differential SCRs. First, the intensive extinction training on Day 2 (96 trials: 48 CS+, 48 CS−) likely attenuated physiological fear responses to the CS+. Second, stimulus novelty effects may have played a role: despite prior exposure on Day 1, the single presentation of perceptually distinct CS+ and CS− images after a 48-hour interval may have elicited novelty-driven orienting responses. Given SCR’s sensitivity to novelty (Zimmer & Richter, 2023), such responses could have masked differential fear reactivity. Finally, the unsignaled shocks administered during reinstatement induced substantial arousal, likely obscuring CS-specific differentiation. Thus, the null SCR findings likely reflect methodological constraints rather than an absence of II effects. Critically, comparable SCR magnitudes between groups confirm that the anger-countering IIs strategy did not elevate participants’ physiological arousal. From a clinical perspective, the absence of increased arousal is beneficial. It aligns with the intervention’s design principles of being “controllable” and “non-cathartic”, demonstrating that utilizing anger as a tool did not introduce unintended physiological stress, impose additional burden or risk, or create compensatory side effects. This stability in physiological state likely supports overall homeostasis and enhances the clinical safety profile of the approach. From a research perspective, however, the failure of SCR to reflect the group differences observed in subjective reports highlights a significant methodological challenge: Validating the successful induction of the targeted state—namely, a transient, conditionally triggered, goal-directed, and controllable form of anger—using peripheral physiological measures like SCR is extremely difficult. The core issue lies in the shared neurophysiological underpinnings of fear and anger: both robustly activate the sympathetic nervous system (Damasio & Carvalho, 2013) and elicit substantial overlap in peripheral physiological responses (Stemmler et al., 2001). This physiological similarity makes it nearly impossible to distinguish arousal caused by fear from arousal specifically triggered by the experimentally induced anger state designed in this study. Consequently, the current research primarily relied on participants’ verbal reports to confirm their subjective experience of anger.

Although the II incorporated both reappraisal and anger induction, our results indicate that cognitive reappraisal was the dominant mechanism. Fear and anger ratings in the II group had already decreased to relatively low levels by the end of the first extinction block, consistent with rapid threat reappraisal (“it is foolish to fear a harmless picture”). By Day 3, the II group also reported lower anger than the control group, suggesting that anger served as a temporary facilitator rather than a sustained emotional state. From a functional perspective, anger was not employed as an uncontrolled cathartic release but as a targeted, conditional, and self-focused response. This design minimized risks of maladaptive aggression and channeled the motivational energy of anger into strengthening reappraisal. Once the irrationality of fear was recognized, anger dissipated naturally. Therefore, the anger in this paradigm was instrumental, transient, and safe, ensuring that approach motivation could be leveraged without escalating hostility. This diverges from prior IIs research where the content primarily focused on cognitive reappraisal devoid of emotional counteraction (Chen et al., 2020, 2021; Gomez et al., 2015; Huang et al., 2020; Ma et al., 2019).

While our data support a cognitive reappraisal-dominated mechanism, the strategic incorporation of anger demands careful consideration. Here, we argue that anger was employed not as a blunt force against fear, but as a targeted, instrumental tool to enhance the cognitive strategy. Its feasibility hinges on its transient and goal-directed nature. Indeed, literature links trait anger and undirected anger to negative outcomes: Research indicates that the association between anger and PTSD may be particularly pronounced in individuals predisposed to impulsive aggression (Teten et al., 2010). Trait anger is a recognized risk factor for developing PTSD (McHugh et al., 2012) and mediates the relationship between PTSD and aggressive behavior (Bhardwaj et al., 2019). Furthermore, theoretical models propose that activation of threat-related cognitive networks can powerfully amplify anger through a positive feedback loop. When this heightened anger combines with combat-related PTSD symptoms characterized by hyperarousal and reactivity, inhibitory control over aggression may be overridden, increasing the risk of violent outbursts (Novaco & Chemtob, 2015). However, the anger elicited by our IIs is fundamentally different. It was conditional (only upon fear detection), self-focused (not outwardly aggressive), and goal-specified (aimed solely at overcoming the fear response). This design ensures that the anger is functional and self-limiting. The anger induction was designed to be non-cathartic. By directing anger inward at one’s own irrational fear, the motivational energy of this instrumental anger was channeled into corrective cognitive reappraisal, rather than an aggressive behavioral release. Once the reappraisal was successful, the anger subsided naturally, preventing it from persisting or escalating. Therefore, the experimental use of anger in this paradigm was not indiscriminate but was highly targeted and conditional. From a mechanistic perspective, these safety parameters (self-focused, non-cathartic) were crucial. They allowed us to test whether anger’s approach motivation could be harnessed without eliciting maladaptive aggression. It may be most suitable for individuals whose primary barrier is high avoidance and low agency, but contraindicated for those with pre-existing anger management issues or impulsive aggression.

A key contribution of this study is evidence that II effects require repeated practice during extinction to become automatized. Prior research has demonstrated that forming IIs enables individuals to automatically regulate negative emotions in anticipated situations (Azbel-Jackson et al., 2016; Chen et al., 2020, 2021; Gallo et al., 2009; Gomez et al., 2015; Hallam et al., 2015; Huang et al., 2020; Ma et al., 2019). However, in these studies, there was typically minimal or no temporal interval between IIs formation and application, and emotion regulation was assessed across numerous trials, with outcomes often reflecting average effects over these multiple applications. Consequently, it remains unclear when and how IIs automate emotion regulation. Specifically, it is unknown whether this automation occurs during initial IIs formation or, like habits, develops through repeated practice. Furthermore, the delayed efficacy of IIs has received limited investigation. In this study, participants formed IIs immediately following fear acquisition on Day 1. Nevertheless, subjective assessments (fear, anger, CS-US contingency, and avoidance distance) conducted prior to extinction on Day 2 revealed no significant differences between the II group and the Ctrl group. This indicates that the IIs formed the previous day had not yet exerted a measurable influence at this early stage, despite participants generally recalling the IIs content accurately. Further, throughout the extinction phase (comprising 3 blocks of 24 trials each), no between-group differences emerged on any measures during the first two blocks. It was only at the conclusion of the final block (Block 3) that a significant reduction in contingency and avoidance distance was observed in the II group compared to the Ctrl group. This pattern suggests that the effectiveness of the IIs required repeated practice during extinction to manifest. Critically, although no IIs instruction on Day 3, the II group exhibited significantly lower levels of fear, anger, avoidance distance, and US expectancy during the reinstatement test compared to the Ctrl group. This demonstrates that repeated practice of the IIs during extinction training on Day 2 led to the automatization of the emotion regulation process, resulting in robust delayed effects that mitigated the return of fear. Integrating the findings from Day 2 and Day 3, it appears that IIs do not automatically trigger the target response upon their formation. Instead, a period of repeated practice is required to transition the execution of the specified emotion regulation strategy into an automated process. Our finding that the IIs’ effects emerged only after practice during extinction suggests that the initial execution of the specified plan requires conscious cognitive effort before it becomes automatized. Furthermore, the automation speed of the responses specified by the IIs may vary across different tasks or behaviors. The surprise recognition test conducted on Day 3 (see Fig. **3F**) revealed no significant inter-group differences in memory performance, which can be explained by two factors: first, the presentation duration of the CS was as long as 5 seconds; second, the execution of the responses specified by the IIs in this study may require very little cognitive effort.

Previous studies have used two main methods to measure avoidance behaviors: (1) No/low-cost avoidance tasks, where participants prevent the US by pressing a simple key (Klein et al., 2021) or moving a joystick (Glogan et al., 2023) during CS presentation; (2) Approach-avoidance tasks (AAT), where choosing CS+ risks shock but yields rewards (e.g., points/money), while choosing CS− avoids shock but incurs losses (Berg et al., 2021; Pittig, 2019; Pittig et al., 2021). Both paradigms have limitations. No/low-cost tasks cause persistent avoidance even when unnecessary (De Kleine et al., 2023; Pittig & Wong, 2021). Costly AATs involve motivational conflicts and cost-benefit trade-offs: while shock intensity is individually calibrated for humans, reward magnitude is often identical across participants, leading to variable avoidance costs (Rattel et al., 2017) and unclear links to extinction outcomes. Additionally, AATs induce a sense of agency (B. Beck et al., 2017) that may reduce fear (Sugawara et al., 2022), and conflate anger- vs. reward-driven approach, complicating result interpretation (e.g., distinguishing II effects from reward contingencies). To address these issues, the present study modified a prior avoidance measure (Aupperle et al., 2011), retaining only the “avoidance” dimension. Participants indicated desired distance from CSs via a 1–9 numerical scale (lower numbers = closer distance/less avoidance), minimizing motivational conflicts for more implicit avoidance assessment.

Several limitations of this study should be acknowledged. First, the sample consisted of non-clinical undergraduates, which limits the generalizability of the findings to clinical populations. Second, the modest sample size (N = 37) and relatively high attrition rate reduced statistical power and may have limited the detection of smaller effects. Third, the use of self-reported anger combined with skin conductance responses (SCR) provided only partial construct validity, as peripheral arousal cannot reliably distinguish fear from anger. Future research should therefore extend this paradigm to clinical populations with fear- and anxiety-related disorders, incorporate larger and more diverse samples, and adopt multimodal physiological and neural measures to better capture the dynamic interplay of fear and anger. Longitudinal designs are also needed to test whether the observed benefits of reappraisal-plus-anger IIs persist over time and across contexts. By addressing these limitations, future studies can more fully evaluate the clinical potential of strategically designed IIs in reducing avoidance, enhancing extinction learning, and preventing relapse in fear-related disorders.

## 5. Conclusion

This study demonstrates that forming and repeatedly practicing implementation intentions (IIs) that integrate cognitive reappraisal with self-directed anger can enhance extinction learning and reduce the return of fear. The primary mechanism appears to be rapid reappraisal of threat, with anger serving as a transient, conditional motivator that facilitates approach and prevents avoidance. Importantly, the strategy required repeated practice during extinction to become automatized, underscoring the role of rehearsal in embedding II-based regulation. By showing that anger, when safely constrained, can be harnessed to strengthen reappraisal, this work extends the Incompatible Response Hypothesis and advances our understanding of how emotion-specific motivations shape fear regulation. Clinically, the findings suggest a novel pathway for augmenting exposure therapy: transforming fear into self-directed anger to increase agency, approach motivation, and safety learning. Future research should test the safety and efficacy of this approach in clinical populations, integrate multimodal physiological and neural measures, and evaluate long-term outcomes. Taken together, these findings highlight the potential of strategically designed IIs to reduce avoidance and relapse in fear-related disorders, offering both theoretical and translational value for the science of emotion regulation.

## CRediT authorship contribution statement

**Hongbo Wang:** Conceptualization, Methodology, Formal analysis, Writing – original draft, review, & editing, Project administration, Funding acquisition, Data curation and Supervision. **Yingzhu Zeng:** Methodology, Investigation, Data Collection, Writing –review, & editing. **Siwen Zeng**: Investigation, Data Collection and Collation.

## Data Availability

Data will be made available on request.

## Acknowledgements

This work was supported by a grant from Ministry of Education, Humanities and Social Sciences Project (20YJC190019), the Science and Technique Foundation in Henan Province (222102310583), the Educational Science Planning Project of Henan Province (2022YB0037).

